# Perception - action dissociations depend on factors that affect multisensory processing

**DOI:** 10.1101/2024.03.24.586486

**Authors:** Nicola Bruno, Stefano Uccelli

## Abstract

Behavioral perception-action dissociations are widely used to test models of high-level vision, but debates concerning their interpretation have underestimated the role of multisensory mechanisms in such tests. Sensorimotor tasks engage multisensory processing in fundamentally different ways in comparison to perceptual tasks, and these differences can modulate the effects of illusion in specific ways in accord with the features of the experimental task. To test this idea, we compared perception and action using a well-understood size-contrast effect, the Uznadze illusion, and manipulated both unimodal and crossmodal stimulation as well as conditions that are known to favor or hinder multisensory integration. Results demonstrate that varying such conditions can cause a visual task to be affected by the illusion, or remain fully unaffected, whereas a visuomotor task can be affected by the illusion, remain immune from the illusion, or, unexpectedly, even show a robust reverse effect. Thus, similar or dissociable effects on perception and action can be observed depending on factors that are known to affect multisensory processing.These findings provide a novel perspective on a long standing debate in behavioral cognitive neuroscience.

## 1. Introduction

It is generally accepted that the visual centers of primate cortex can be grouped into two systems [1], roughly corresponding to areas located below (“ventral”) and above (“dorsal”) the Sylvian fissure. Based on neuropsychological and neuroimaging data, these two systems have been proposed to reflect a functional distinction between processing of object-relative information about shape and color, for the purpose of recognition and classification, and processing of observer-relative information about location and movement, for the purpose of controlling action [2]. Within this framework, behavioral dissociations between perceptual and motor responses to visual illusions are widely cited as key evidence for the functional independence of the two systems [3]. However, the empirical status of behavioral perception-action dissociations in perceptual neuroscience remains controversial to this date [4-6].

For instance, a seminal study reported that the maximum in-flight aperture of the digits when grasping a test object remains proportional to its physical size, despite perceptual size-contrast effects due to flanker objects [7]. However, a large-scale pre-registered multicenter study reported similar effects of size-contrasting flankers on both finger apertures in grasping and on three different measures of consciously perceived size [8], and a recent study found similar effects on perception and action from larger (or smaller) objects (size contrast “inducers”) presented immediately before a test object [9]. While several explanations have been proposed, there is currently no consensus on what causes these inconsistent results.

A neglected issue in the interpretation of perception-action dissociations concerns the multisensory nature of human perception [10]. Although it is generally believed that visuomotor tasks only engage visual coding, these protocols in fact engage not only visual but also somatosensory processes [11-13], and they do so in fundamentally different ways in comparison to perceptual tasks. To be true, some studies did pay some attention to the potential impact of haptic information in perception-action comparisons, but focussed merely on the potential adapting effect of obtaining haptic feedback, once the grasp has been completed (e.g., [14-15]). We argue that these potential effects, while relevant, did not address a key issue for perception-action studies. This is so because, while they could introduce uncontrolled differences between perception and grasping tasks, adaption effects such as these can be readily eliminated by modifying experimental procedures. For instance, in many studies (e.g. [8]) that involved comparing grasping to matching, participants were required to pick up the test object after matching to equate haptic feedback in the two conditions.

Much more fundamental multisensory effects are however likely to play a role already at the planning stage of motor responses.

Consider grasping an object, which is arguably the most studied visuomotor task in this literature. It is well-known that the motor representation of object size, as indexed by the maximum in-flight aperture of the digits as the hand approaches the object, is a monotonic function of the actual size of the object [16]. This-well-established finding suggests that visuomotor processing accesses representations of object size when planning grasps. This finding also implies, however, that visuomotor processing also accesses representations of one’s body. Specifically, such processing will need to access representations of the hand’s three-dimensional structure as the digits will need to open proportionally less with bigger hands than with smaller ones to achieve the observed scaling. Thus, the representation of object features for planning grasps will necessarily recruit multisensory processing of visual and somatosensory mechanisms.

Supporting this idea, there is evidence that experimentally altering the internal representation of one’s hand [17] will modulate the maximum grip aperture while grasping [18-20]. In addition, there is evidence [21] that when repeatedly grasping an object that (haptically) feels larger (or smaller) than it looks, the maximum grip aperture correspondingly also becomes larger (or smaller). Although this effect has been described as a form of multisensory adaptation, a parsimonious explanation is again that the internal representation of the hand was modified by the contrast with the felt object, such that repeatedly feeling a large object caused the hand to feel smaller (or vice versa). Note that, perceived object size being the same, grasping with a smaller hand requires opening the fingers more, while grasping with a larger hand will tolerate a smaller aperture.

In contrast, consider reporting the perceived size of the same object. Techniques to assess such reports typically consist of matching tasks, which can be broadly grouped into unimodal and crossmodal. In unimodal matching, participants report perceived size by verbally indicating which comparison item best corresponds in seen size to a displayed test object. Under the assumption that language-related motor processes to report the match remain encapsulated from sensory processing [22], this task can be reasonably classed as purely visual. In crossmodal matching (often also dubbed “manual estimation”), participants instead report perceived size by matching it to the aperture of the index and thumb fingers of one hand. This type of match necessarily requires comparing a visual signal, the seen size of the object, to a somatosensory signal, the felt distance between the digits. There is evidence that perturbations of such somatosensory signal, again by altering the internal model of the hand, can modulate haptically felt size, but this effect is in the opposite direction to that on the grasping maximum aperture [23] (see Discussion).

Here we compared perception and action in conditions involving both visual and haptic stimulation and that are known to favor or hinder multisensory integration. To this aim, we exploited the Uznadze illusion [24] (Fig. 1a), a well-understood size-contrast effect whereby a larger (or smaller) inducing stimulus causes a test object to appear smaller (or larger). The Uznadze illusion is ideally suited to explore multisensory effects on perception and action. The illusion effect is large (10-20% of increment/decrement) and readily observable in most participants [24-25], which greatly reduces the uncertainty of parameter estimates in statistical comparisons. In addition, equivalent effects are observed both in haptic [24][26], in visual [25] [9] [27], and audiomotor [28] versions. In each trial of our experiment (Figure 1c and 1d), participants were first exposed to an initial inducer stimulus which could be larger, smaller, or the same size as a test object that participants were required to grasp after exposure to the inducer. The inducer stimulus could be either visual or haptic, and it could be presented in the same (frontal) or in a different (lateral) location relative to the frontal grasping test. After the grasp, which was always performed with the right hand, participants reported the perceived size of the test by matching it to the aperture of the index and thumb of the left hand. Thus, the grasp was always visually-driven, but could be affected by the inducer either unimodally (when the inducer was also visual) or crossmodally (when the inducer was haptic). Conversely, the match always entailed a unimodal haptic comparison.

**Figure 1.**
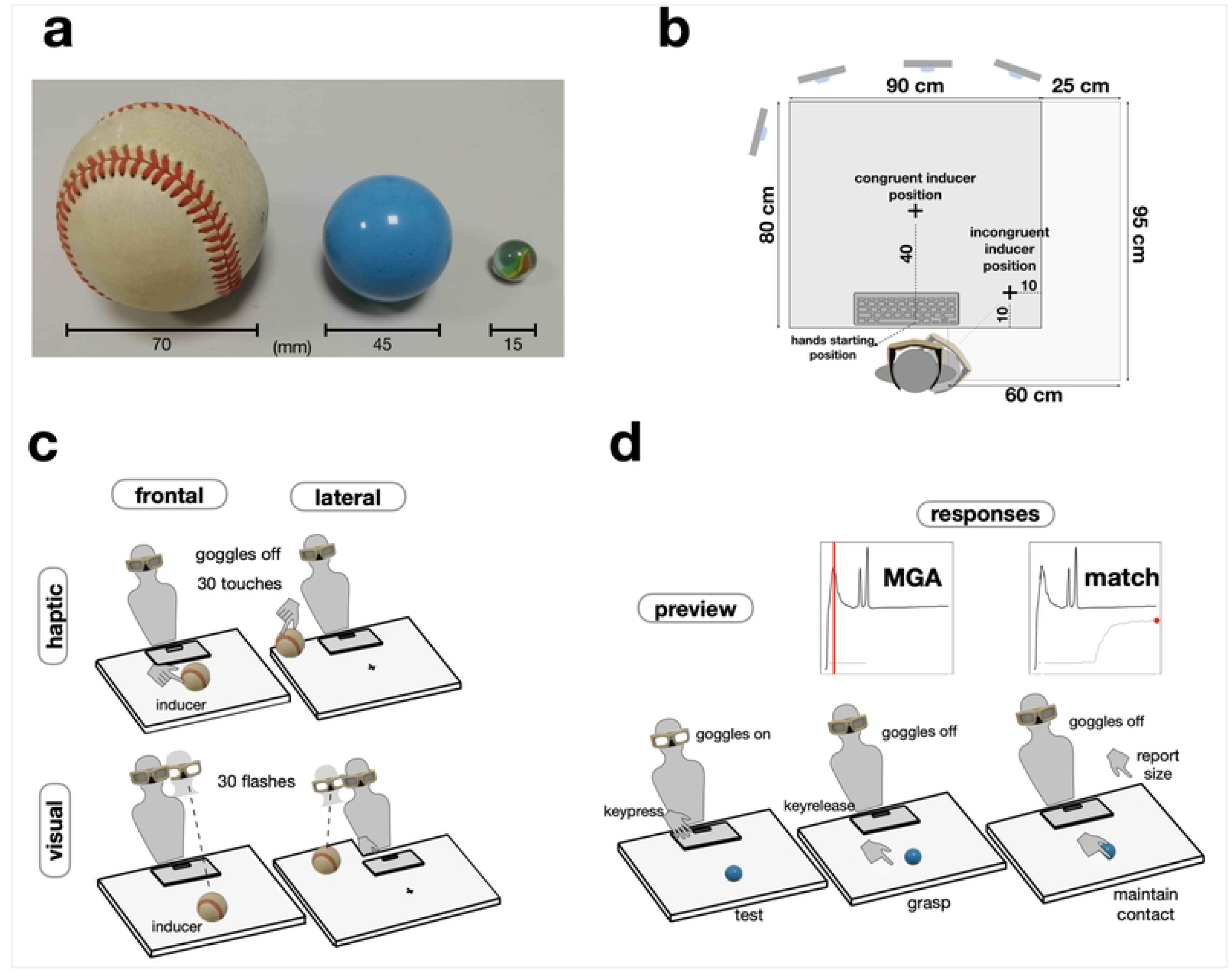
Overview of methods. **a**. Stimuli were three commercially available spherical objects, a regulation baseball, a rubber ball, and a glass marble. All three were used as inducers, whereas the test object was always the rubber ball. Thus the test could be either larger or smaller than the inducer, or the same (baseline) as the inducer. In the Uznadze illusion, exposure to a smaller inducer causes the test to appear larger than baseline, and exposure to a larger inducer has the opposite effect. The illusion is readily observed with haptic [24] visual [25], as well as visuomotor [9] measures. The difference is typically 10-20% of physical size. **b**. Participants sat in front of a square table with a computer keyboard on its near edge and four recording cameras around it. Graspable tests were always placed 40 cm sagitally in alignment with the participant midline. Inducers were placed laterally on the right or sagitally. In visual/lateral blocks, an extension was applied to the table (light grey) to insure that inducers were seen against a homogenous background as in the frontal blocks. **c**. The induction phase of a trial could involve visual or haptic stimuli that could be presented either frontally (spatially congruent with tests) or laterally (incongruent). Spatial congruence is known to be a key factor favoring multisensory integration [30]. **d**. After induction, observer previewed the frontal test, grasped it with the right hand, and then matched its perceived size with the left hand. Both grasping and matching were executed without vision as the shutter googles turned off as soon as participants lifted their hand from home position.

## 2. Methods

### 2.1 Ethics

All methods and experimental protocols were developed in accordance with relevant guidelines and regulations. Specifically, the study was conducted following the ethical standards of the Code of Ethical Principles for Medical Research Involving Human Subjects of the World Medical Association (Declaration of Helsinki), with the ethical standards of the Italian Board of Psychologists (see https://www.psy.it/codice-deontologico-degli-psicologi-italiani), as well as the Ethical Code for Psychological Research of the Italian Psychological Society (see https://aipass.org/node/11560). All experimental protocols have been approved by the Research Ethics Board (REB, 49-2022-N) for non-medical research involving human participants of the University of Parma. As the study did not involve clinical tests and medical equipment, and no health information was collected from participants, verbal consent only was deemed a prerequisite for participating.

Potential participants were informed beforehand that their anonymity would be preserved at all stages. All participants gave informed verbal consent after being adequately informed about the aims, procedure, and methods of the study and were free to abandon the experiment at any moment.

### 2.2 Participants

Ten members of the University of Parma community participated (4 females, mean age: 30.5; range: 24 - 58) in the main study. They were the authors (N.B. and S.U.) and 8 volunteers who were naïve to the aims of the study. The recruitment started on 2nd March 2021 and ended on 10th June 2021. An additional ten participants (7 females, mean age: 25.8; range: 22 -35) participated in a second study involving only the “haptic induction” conditions (see section 2.6). The recruitment started on 8th February 2022 and ended on 13th April 2022.

### 2.3 Sample sizes

The number of participants for the main study was limited to 10 for the reasons detailed below. First, the haptic size-contrast effect used in the study (the Uznadze illusion) is a well-understood, robust phenomenon. Individual differences are negligible and reliable estimates of effect sizes are available for both perceptual [25] and sensorimotor tasks [9]. We therefore expected to observe it in the relevant experimental condition in essentially all participants. Second, the visual induction conditions of the study (see 2.5) mimicked those of our previous study [9] and we expected that they would readily replicate our previous results, as they indeed did. Conversely, we had no specific predictions regarding the novel haptic induction conditions. These conditions yielded an unexpected result and for this reason we performed an exact replication of the study in these conditions only. The results of this replication are reported in 3.2 and they confirm the validity of the expected result. Third, and final, we analyzed not only group data but also individual patterns. This approach has been shown to have superior power and inferential validity in comparison to traditional group comparisons [29].

### 2.4 Stimuli

We used three commonly commercially available balls: A regulation baseball (large, diameter ≈ 70 mm), a rubber toy ball (medium, 45 mm), and a marble (small, 15 mm). Figure 1a displays them. All three were used as inducing stimuli, whereas only the medium ball was used as a test stimulus. Exposure to a large inducer yielded a successive contrast that was expected to reduce the perceived size of the test; exposure to a small inducer yielded a contrast expected to enlarge its perceived size; and exposure to a medium inducer generated a baseline no-contrast condition for comparison.

### 2.5 Task

The experimental task consisted of a sequence of three distinct phases. In the first phase, participants perceptually explored an inducing stimulus (“induction”). In the second phase, they previewed, reached for, and grasped a test stimulus, lifted it, and placed it on the table ∼10 cm to the right of its original position (“grasping”). In the third phase, finally, participants reported on the perceived size of the test by providing a match with the left hand while keeping the test in the right hand (“matching”). In the induction phase, inducers were presented either haptically or visually. In the haptic induction conditions, participants explored the inducing stimulus by moving the right-hand thumb and index fingers around its surface (i.e., a ‘contour following ‘exploratory procedure [31] without visual information (shutter glasses closed). In the visual induction conditions, participant fixated and observed the inducing stimulus (shutter glasses open) with their right hand at home position.

Consistent with previous assessments of the Uznadze illusion [25; 9; 32] the haptic and visual induction phases both lasted 7-8 s (see 2.8). In the grasping phase, participants were instructed to use the index and thumb of the right hand, which was the same hand they used to explore haptic inducers, and to maintain active contact with the test stimulus after having lifted and placed it again on the table. To this aim, they were encouraged to perform repeated finger presses to maintain awareness of the stimulus size. In the matching phase, finally, participants lifted their left hand on their side and adjusted the aperture of the index and thumb fingers to match the size of the test. Both the grasping and the matching responses were performed in visually open-loop conditions by means of shutter glasses. Specifically, participants were allowed to preview the tests as long as they wished, but as soon as they lifted their hand to reach for it, the glasses closed and everything else was done without visual feedback.

### 2.6 Design

In each trial, we recorded the maximum in-flight aperture (*maximum grip aperture*, MGA) as the right finger and the thumb approached the test stimulus during the grasping phase, as a function of three independent variables: *inducer size, spatial congruence*, and *inducer modality*. Specifically, inducers could be either large, small, or medium (that is, the same size as tests). Inducing stimuli were presented either in the same position (in front of the participant at arm’s reach) or in different positions (inducer on the right side and test frontal) as the tests, and they could be either visual or haptic. Each inducer was presented 20 times. For each of the four conditions we run one 60-trial block, such that the experiment totalled 240 trials per participant. The four blocks were counterbalanced across participants over two consecutive days (two blocks per day). Each block was further subdivided into 5 sessions of 12 trials to reduce fatigue. Inducer size was randomized within sessions. Overall, each block required about one hour to complete and the whole experiment therefore lasted about 4 hours.

### 2.7 Apparatus

Participants sat on an adjustable chair in front of a rectangular table (90 × 80 cm). Faint cross-marks drawn on the table marked the position of the inducing and test stimuli. In all conditions, tests were placed on the cross-mark positioned at 40 cm from the spacebar (40 cm from the table’s edge); both the cross-mark and the center of the spacebar were aligned with the participant’s midline. Conversely, positions were different between congruent and incongruent spatial conditions: In the former, inducers were placed on the same cross-mark as the test; in the latter, they were placed on the cross-mark to the right of the participant at 10 cm from the right edge of the table (35 cm from the spacebar). In the visual incongruent condition, the area of the table was extended by adding a larger panel on it. This was done to present inducers against a homogeneous background that was comparable to that of the frontal inducers. Schematics of the experimental apparatus are presented in Figure 1b. The visibility of all stimuli was controlled by PLATO shutter glasses (Translucent Technologies Inc., Toronto, Ontario, Canada), which have a transparent-to-opaque latency of 3-4 ms. Kinematics were recorded using a BTS-DX100 Smart System (sampling rate 100 Hz, spatial precision < 0.2 mm with four cameras at distances of 1-1.5 m from the participant’s hand). Three markers were placed on the wrist, the tip of the index, and the tip of the thumb of the right hand; two additional markers were placed on the tips on the index and thumb of the left hand. A personal computer running MATLAB (2013b, The MathWorks, Natick, MA, USA) under Windows 7 was used to control stimulus presentation and to issue trigger signals to the glasses and the tracking system.

### 2.8 Procedure

Trial structure is schematized in the middle and bottom panels of Figure 1c and d. Each session began with a verbal explanation followed by 6 practice trials with randomly chosen inducers. Participants then performed the experimental trials. At the beginning of each trial, the shutter glasses were turned off (no vision), and the experimenter picked up the appropriate inducer from a lateral container and placed it on the table either sagittally in front of participants or on their right side as appropriate for the current block. At this point, the experimenter gave a go signal indicating to participants to initiate the trial. In visual blocks, this was done by participants by pressing the spacebar of the keyboard with their left hand, which caused the shutter glasses to open (25 ms) and then close (25 ms) for 30 times, mimicking a visual exploration of the inducer. At the end of 30th cycle, the shutter glasses remained closed and this concluded the visual induction phase. In the haptic blocks, trials began with shutter glasses off while participants actively touched the inducer stimulus placed on the right side for 30 times (4 touches/s), such that participants could feel it haptically using the thumb and index fingers of their right hand. At the end of the 30th touch, the shutter glasses remained closed and the induction phase was similarly concluded. At this point, the experimenter removed the inducer and placed the appropriate test in front of participants. A second go signal was then given, indicating to participants that they had to press the spacebar in preparation for grasping. This second spacebar press opened the shutter glasses, which remained opened until the participant released the spacebar to initiate the grasp and to start recording the kinematics of the response. At this point, the participant grasped the test stimulus, lifted it, and repositioned it on the table. This concluded the grasp phase of the trial. In the final, matching phase, the participant lifted the left hand and reported the perceived size of the test by opening the thumb and index. The left hand index-thumb aperture was maintained until an acoustic signal informed that the trial (and the kinematics recording) ended. The recording time from the spacebar release to the end trial signal lasted 10 s, giving participants ample time to perform the whole sequence.

### 2.9 Data validation

The kinematics of all trials were screened for atypical grasping profiles. Trials that met the following criteria were included in further analyses: 1) for the right hand, plotting the distance between the thumb and index as a function of time revealed an initial unambiguous peak (the MGA); 2) for the left hand, the corresponding distance evolved rapidly and then remained approximately constant until the end of the recording period. Trials classified as incorrect were discarded. According to these criteria, data validation resulted in the exclusion of only 13 datapoints (∼ 0.5%) based on criterion 1 and of 12 datapoints (∼ 0.5%) based on criterion 2. Both the MGA and the match were computed from kinematics records by means of custom-made R scripts, by inspecting the vector of the index-thumb distances (see red vertical lines in Figure 1d). The qualitative pattern of the data was first evaluated by plotting MGAs and matches as a function of inducer size individually for each participant in each condition (Figure 2).

**Figure 2.**
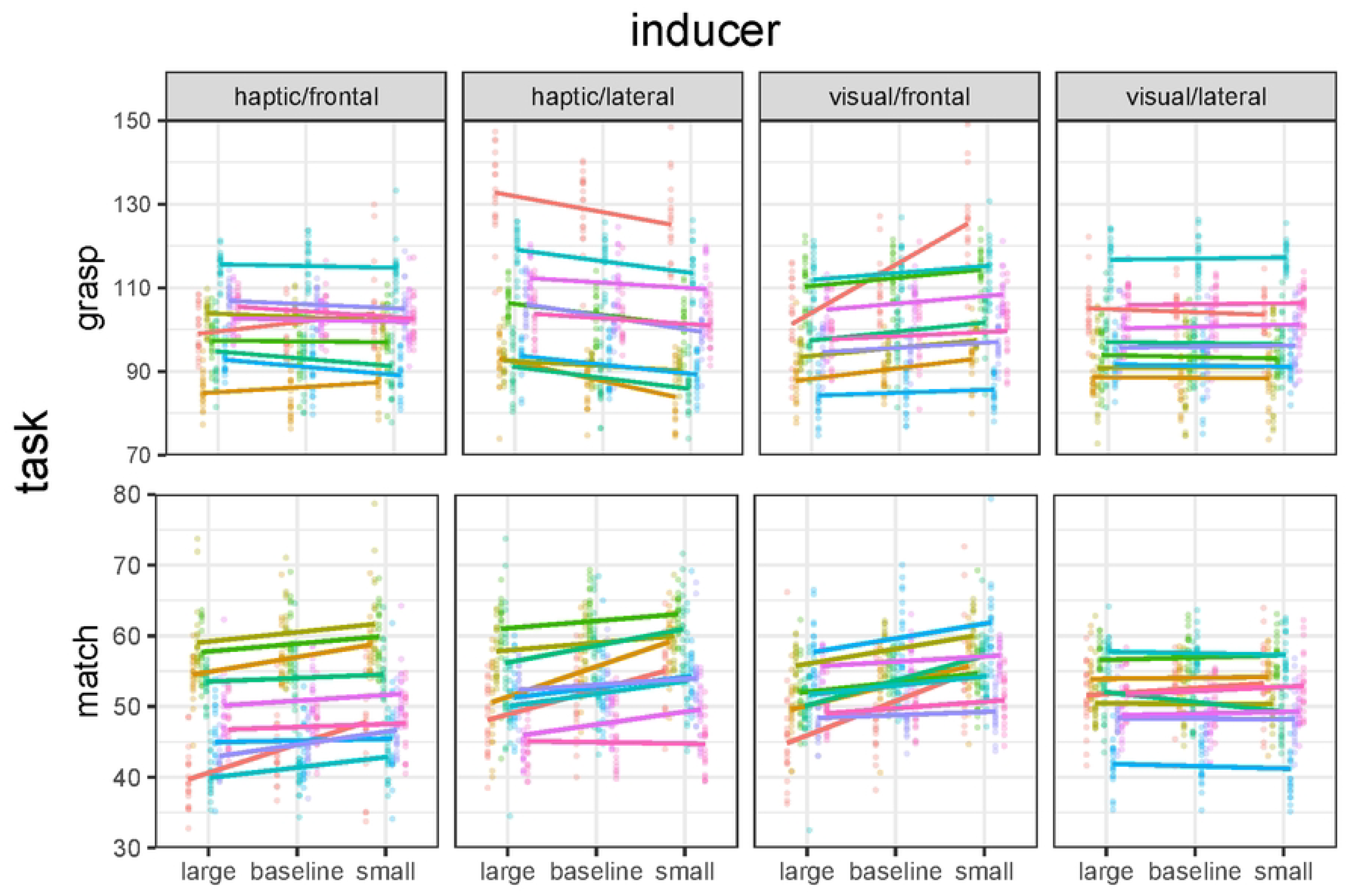
Individual finger aperture data in grasping and matching. Columns: conditions (frontal inducers were spatially congruent with tests, lateral inducers were not). Grasp data: maximum in-flight aperture (MGA). Match data: haptic size match (“manual estimation”). Points: individual distributions at each size level. Segments: linear regression fits to the individual data. Colors identify participants.

### 2.10 Outlier detection

Visual inspection of the plot in Figure 2 revealed extreme values in both measures across participants and conditions. Accordingly, before proceeding with further analyses we screened the data for outliers by detecting values beyond ± 2.5 in the standardized residuals individual distribution of the linear model fitting the data (Figure 3). Accordingly, the 1.8 and 2 % of the data was excluded from the grasp and match data, respectively.

**Figure 3.**
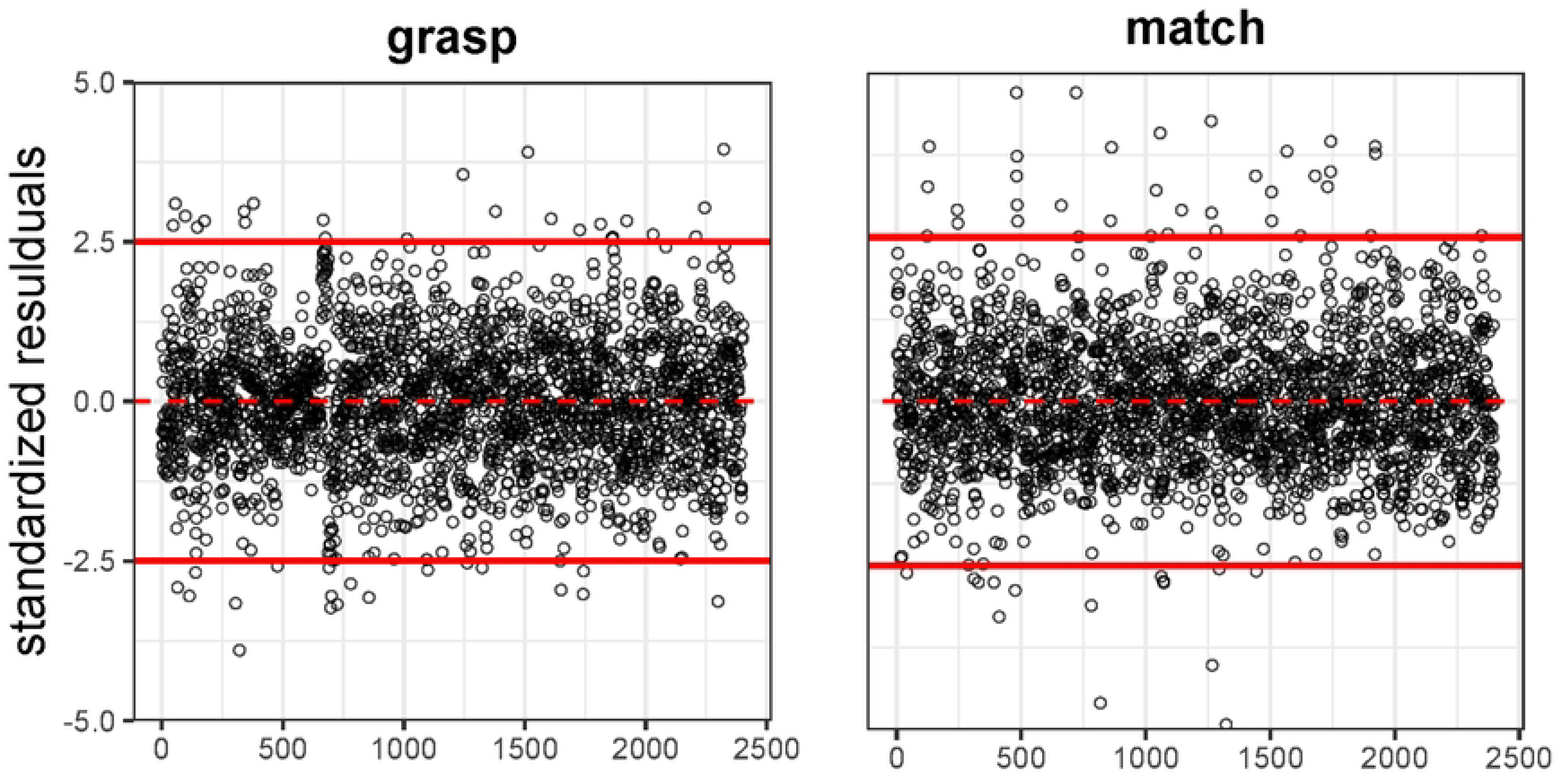
Residual plots. Residuals from the linear model fitting the individual grasp and match data as a function of inducer size and experimental condition were inspected to detect outliers.

### 2.11 Re-expression

Individual differences were observed in the average value of both intercepts and slopes of the linear model fitting individual datapoints. Given that differences between intercepts are irrelevant to the current experimental question, we re-expressed both measures by subtracting individual averages in the baseline conditions from each datapoint (see [33-34]). This yielded a new plot which represented inducer effects as deviations from the baseline average, with negative values signalling a shrinking effect and positive values signalling an enlarging effect of inducers on tests (i.e., the Uznadze effect, Figure 4). We used these re-expressed data to model our results using individual slopes (which measure the Uznadze effect) as random factors, and to estimate confidence intervals (CIs) on parameter estimates.

**Figure 4.**
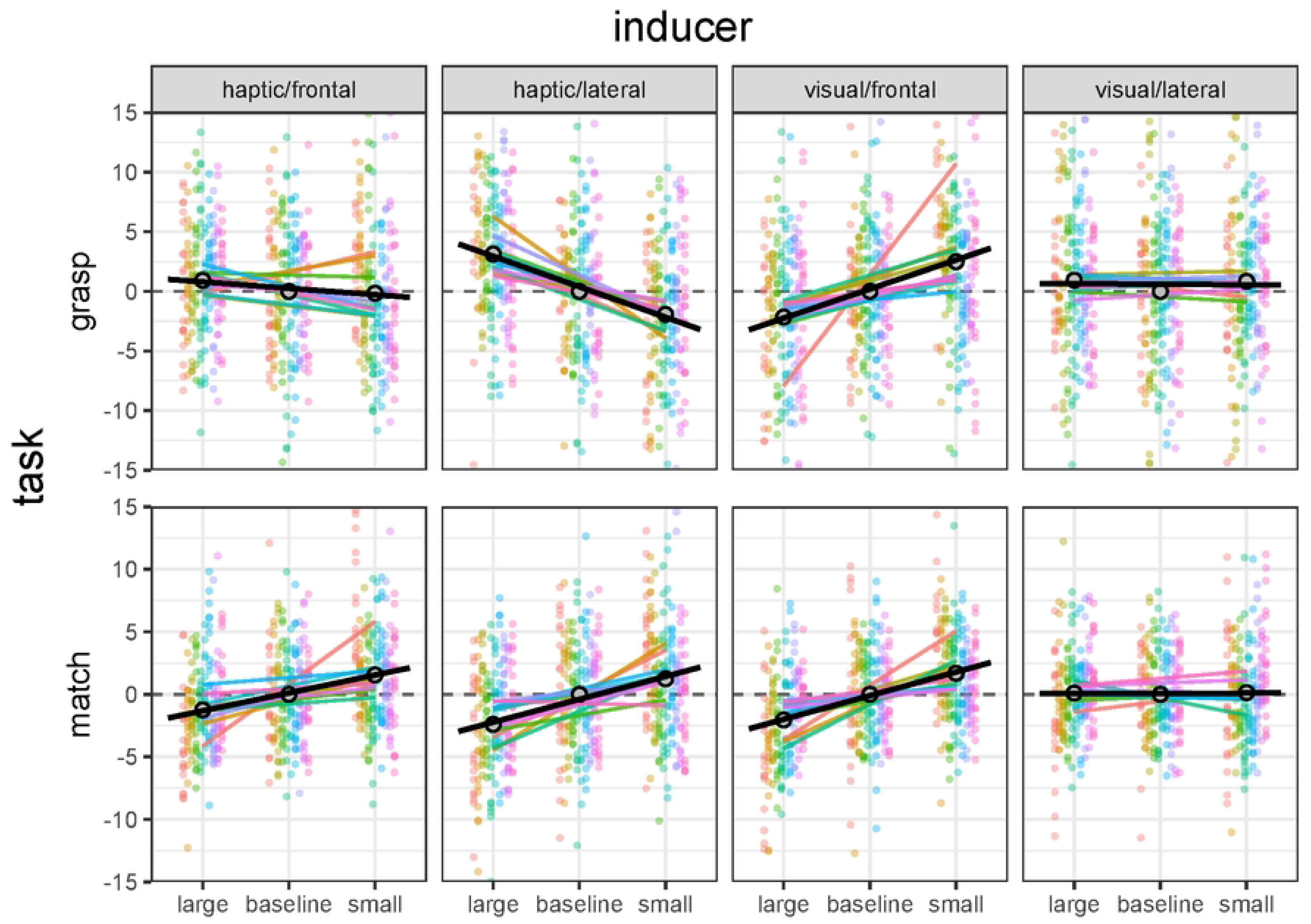
Individual finger aperture data re-expressed as deviations from the baseline average in the grasping and matching phases of our task. Each data point is a single trial from one participant, thin lines are regression fits to the individual data, colors identify participants. Thick black lines are the average regression fits obtained by averaging individual intercepts and slopes.

### 2.12 Statistical approach and Analysis

The re-expressed grasping and matching data were modelled with linear mixed-models (LMMs) [35] in R (4.2.2) using the function *lmer* from the *lme4* package [36]. In contrast to traditional analysis of variance, which uses ordinary least-squares, LMM uses maximum-likelihood estimation to compute likelihood-ratios (λ_LR_). This has several advantages. First, it allows to predict participant-by-participant variation in model parameters (random effects) and to discount these individual differences to compute within-participant CIs (95%) around estimates of fixed effects (i.e., the experimental conditions). Second, LMM uses the Akaike information criterion (AIC, see [37]) that allows to compare models having different numbers of parameters.

Third, LMM allows comparisons across conditions based on CIs around estimates instead of traditional p-value tests, in accord with current recommendations on analytical approaches in psychology and behavioural neuroscience [38-40].

## 3. Results

### 3.1 Main study

A summary view of results from our main study is presented in Figure 5, which reveals a complex pattern of dissociations between perception and action. In perception, the Uznadze illusion appears to affect matches in all conditions except when the inducer was presented laterally and the test was frontal (visual/lateral condition). In action, the illusion seems to affect the MGA in the visual/frontal condition but not in the visual/lateral and the haptic/frontal conditions. Surprisingly, the illusion appears to affect the MGA also in the haptic/lateral condition but in this case the effect is reversed: participants opened their fingers more after exposure to a large inducer, and less after exposure to the small inducer.

**Figure 5.**
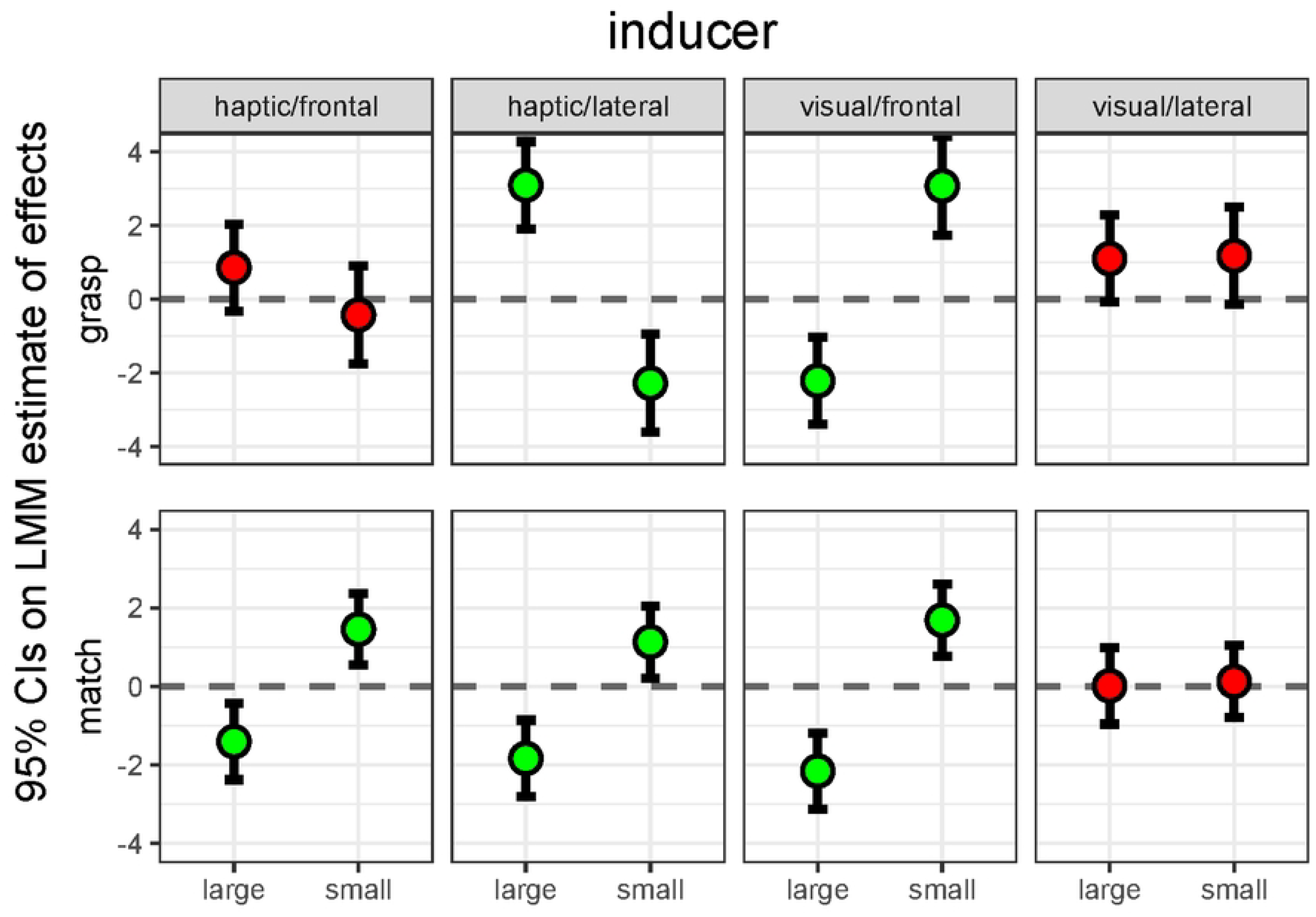
Effects of the visual and haptic Uznadze illusion on the visuomotor (grasp) and perceptual (match) coding of object size. Plots depict 95% confidence intervals on the estimated effects of large or small inducers on grasp and match finger apertures. Effect estimates and associated standard errors were derived from fitting a linear mixed model (see Results). The reference horizontal line at y = 0 represents the intercept of the model (i.e., baseline data), and plots in green identify conditions whereby we are 95% confident that there is a difference between the effects of the two inducers. The results reveal a complex pattern of dissociations between and within the perception and action tasks. In the matching data, inducers affected perceived size in all conditions except when a visual inducer was presented laterally and the test was frontal (visual/lateral condition). In the grasp data, inducers did not affect visuomotor size in the visual/lateral and the haptic/frontal conditions, whereas they affected visuomotor size in the visual/frontal and haptic/lateral conditions. However, in the former the effect was consistent with the classic Uznadze effect (shrinking after a large inducer and expansion after a small inducer), whereas in the latter it provided evidence for an inverse Uznadze (shrinking after a small inducer and expansion after a large one). Thus, inducers caused size contrast, size assimilation, or no effect, depending on the conditions of exposure.

This qualitative assessment of results was supported by estimates of effects and of their confidence intervals. Confidence intervals were estimated based on model selection after comparing different linear mixed models (see following paragraph). Specifically, for both the matching and grasping data CIs were based on estimates from the model predicting finger apertures from the interaction of the inducer size and the experimental condition. Figure 5 presents these effects with 95% CI as a function of these two factors. Comparisons can be performed by noting distances between upper and lower CI limits in relation to the horizontal dashed lines (zero effect expectation). Table 1 in the online supplemental materials presents the exact numerical values of estimates.

Modelling was performed as follows. Participants were treated as the random factor. Inducer size and the experimental conditions were treated as fixed factors. The former consisted of three-levels, while the latter was treated as a unique four-level factor. Using a maximum likelihood criterion we compared the fit of a generalized least squares (GLS) null model with fixed intercept and slope (1) with that of a null model with random slope (2). Then, the factor ‘experimental condition’ was added as a fixed effect, generating model (3) for comparison to model (2). Next, the factor ‘inducer size’ was added as a fixed effect, generating model (4) for comparison to model (3). Last, we modelled the modulation of the inducer size effect by the different experimental conditions. This was done by entering the size X condition interaction as a fixed effect generating model (5). In the matching data, model (2) revealed a better fit than model (1) [ΔAIC = -154; λ_LR_: χ^2^(8) = 165.9, p < .001]. Model (3) did not reveal a better fit than model (2) [ΔAIC = 4; λ_LR_: χ^2^(11) = 0, p = 1]. Model (4) revealed a better fit than model (3) [ΔAIC = -9; λ_LR_: χ^2^(10) = 12.3, p = .002]. Model (5), finally, revealed a clear better fit than model (4) [ΔAIC = -35; λ_LR_: χ^2^(19) = 63.3, p < .0001]. In the MGA data, model (2) did not reveal a better fit than model (1) [ΔAIC = 6; λ_LR_: χ^2^(8) = 5.9, p = .42]. Model (3) did not reveal a better fit than model (2) [ΔAIC = 3; λ_LR_: χ^2^(11) = 0, p = 1]. Model (4) did not reveal a better fit than model (3) [ΔAIC = -1; λ_LR_: χ^2^(10) = 5.3, p = .07]. Model (5), finally, revealed a clear better fit than model (4) [ΔAIC = -142; λ_LR_: χ^2^(19) = 161.3, p < .0001]. Thus, LMM indicated that in both the MGA and matching data outcomes were affected by the size of inducers but this effect, crucially, was modulated by experimental conditions.

### 3.2 Replication of haptic inducer conditions

Given the unexpected results observed in the haptic inducer conditions (inversion of the illusory effect), we performed an exact replication of the relevant conditions (haptic inducer, lateral vs haptic inducer, frontal). The replication was performed to rule out unlikely, but not impossible, distortions due to the small number of participants employed in the main study. All methods were exactly the same as before, including data validation and data analysis. A summary of results are presented in Figure 6. Table 2 in the online supplemental materials presents the exact numerical values of estimates.

**Figure 6.**
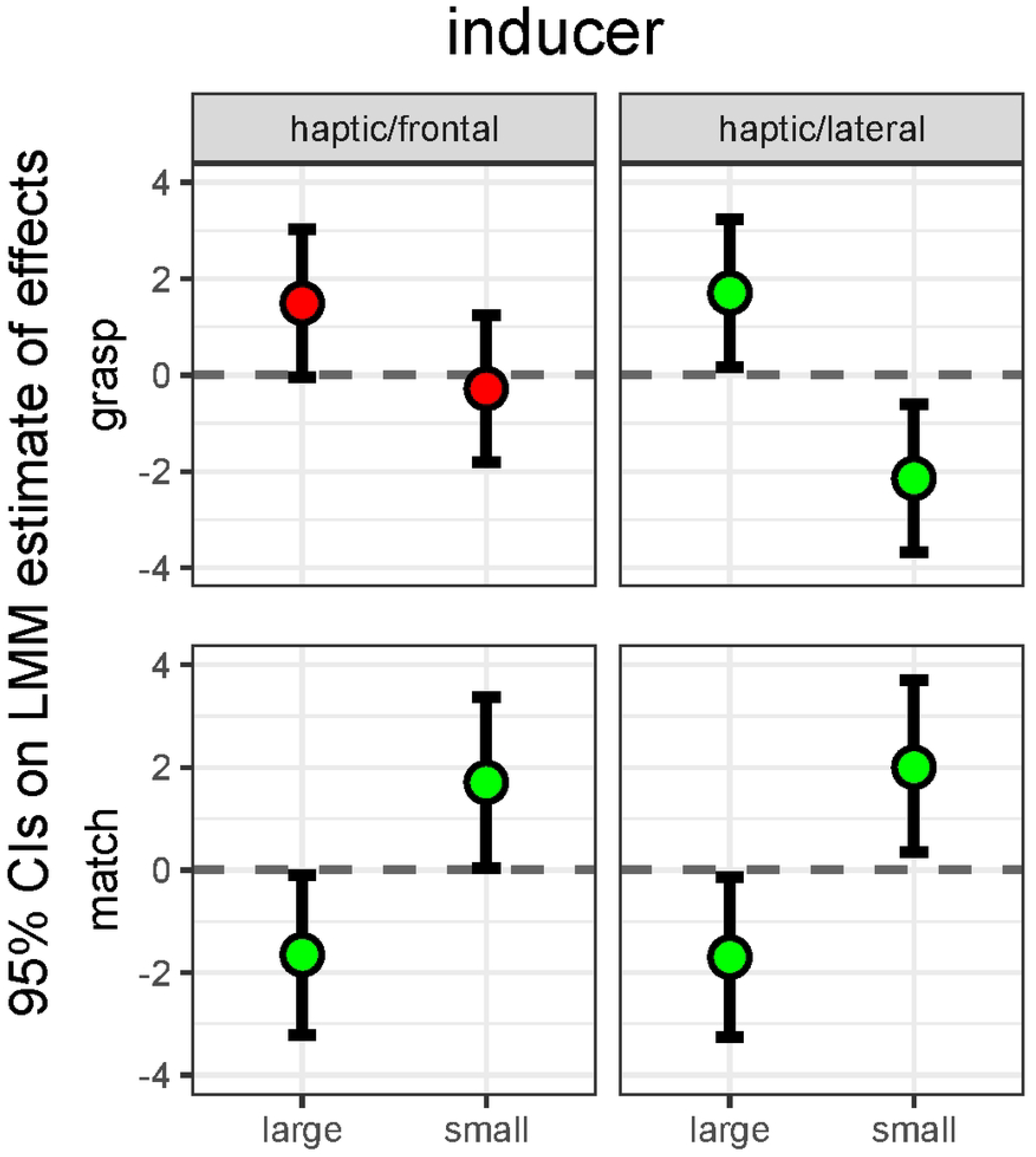
Replication of the haptic inducer conditions from the main study. All plotting conventions as in Figure 5.

Results closely resemble those of the main study. Modelling these data also yielded very similar results to those of the main study. In the matching data, model (2) revealed a better fit than model (1) [ΔAIC = -180; λ_LR_: χ^2^(8) = 191.7, p < .001]. Model (3) did not reveal a better fit than model (2) [ΔAIC = 1.9; λ_LR_: χ^2^(9) = 0.09, p = .76]. Model (4) revealed a better fit than model (3) [ΔAIC = -3.9; λ_LR_: χ^2^(10) = 5.9, p = .01]. Model (5), finally, did not reveal a better fit than model (4) [ΔAIC = 5.6; λ_LR_: χ^2^(13) = 0.39, p = .9]. In the MGA data, model (2) revealed a better fit than model (1) [ΔAIC = -19; λ_LR_: χ^2^(8) = 31.3, p < .001]. Model (3) did not reveal a better fit than model (2) [ΔAIC = 0; λ_LR_: χ^2^(9) = 2, p = .15]. Model (4) revealed a better fit than model (3) [ΔAIC = -16; λ_LR_: χ^2^(10) = 17.8, p < .0001]. Model (5), finally, revealed a slightly better fit than model (4) [ΔAIC = -2.1; λ_LR_: χ^2^(13) = 8.15, p = .04]. Given that these results are essentially identical to those from the main study, we are confident that the haptic inducer results are not due to distorted estimates in a small sample.

## 4. Discussion

Consider first the visual inducers (Figure 5, third and fourth columns). When these were spatially congruent with the tests, we observed equivalently narrower or wider apertures in grasps and in matches. In contrast, when visual inducers were not congruent, they had no detectable effect on either the grasps or matches. Thus, the Uznadze illusion produced similar effects on both perception and action, as already reported in an earlier study of our group [9]. This is consistent with an involvement of multisensory processes in the programming of the grasp. When inducers and tests were spatially congruent, visual information about inducers appears to interact with the somatosensory and sensorimotor processes involved in grasping the tests. When inducers and test we not congruent, the visual coding of inducers instead appears not to interact with the coding of the test either in sensorimotor processes (when grasping) or in the later haptic coding (when matching).

It has been long known that the integration of separate sensory signals, such as vision and haptics, mostly tends to occur when such signals originate from coincident spatial locations. In non-human animals, this fundamental principle of multisensory perception has found support from electrophysiological recording of neurons in the superior culliculus [41-42]. However, it is generally believed that the same principle applies to perceptual responses from intact behaving organisms [43], at least in tasks that require spatial coding [44]. There is also evidence that nonspecific multisensory processes, for instance in adaptation paradigm, can affect grasping kinematics [45-46]. Our findings in the visual inducer conditions are therefore consistent with an engagement of multisensory processing on both the grasping and the matchings tasks, as one would expect from known principles of multisensory integration.

Now consider the haptic inducer conditions (Figure 5, first and second columns, Figure 6). Here, the pattern of the sensorimotor and perceptual effects was surprisingly different from that observed with visual inducers. Independent of whether they were lateral (and therefore felt in a different location than the later grasping test) or frontal (in the same location), large haptic inducers reduced perceptual matches, whereas small inducers increased them. Thus, the haptic Uznadze illusion affected matches independently of spatial congruence.

Conversely, our estimates of mean inducer effects suggests (up to 95% confidence) that inducers did not affect sensorimotor apertures when they were congruent with the tests (frontal), but affected sensorimotor apertures when they were not (lateral). This pattern however was the inverse of the classic Uznadze illusion: Large inducers yielded larger, not smaller, in-flight apertures, whereas small inducers yielded smaller apertures.

This complex pattern of results is again predicted by the role of spatial congruence in multisensory integration, combined with the accessory principle that multisensory experience rapidly alters internal models of the body [10]. There is evidence that such modifications modulate both the in-flight aperture in grasping [18] [20] and perceptual haptic reports [23], but do so in opposite ways. Consider first the effect of haptically exposing participants to large or small inducers on grasping. Exposure to large inducers will tend to shrink the internal representation of one’s hand, and exposure to small inducers will tend to enlarge it. As a consequence of this plastic change in the internal model of the hand, visual size being equal, actors will tend to open the fingers more after exposure to large inducers (the hand is smaller) and less after exposure to small inducers (the hand is larger). This will produce an inverse Uznadze effect, as was indeed in the our data. Now consider the corresponding effect on matching. When matching the test sizes, instead, participants will compare the size of the test, as felt by the digit aperture in their right hand, to the aperture of the digits in the left hand. The former will be calibrated to either a smaller hand (large inducer), yielding a smaller perceived size, or a larger hand (small inducer), yielding a larger perceived size. Assuming that the change in the internal hand model only happens in the exposed hand, one would predict the classic Uznadze effect, as was again the case in our data. Classic data from prismatic adaptation studies support the assumption that sensorimotor recalibration is confined to the adapted hand [47].

Note, however, that these multisensory effects will combine with those of the object-relative size contrast between the inducer and the test when these are spatially congruent. Accordingly, one would expect to observe that in-flight apertures in the frontal condition (spatial congruence of inducer and test) will reflect a compromise between the hand-relative effect, which produces an inverse Uznadze, and the multisensory object-relative counterpart, which produces the classic Uznadze. As these effects are in opposite directions, one would expect that they will tend to cancel out, as is again the case in our data.

## 5. Conclusion

These results provide a novel perspective to the debate on behavioral dissociations between perception and action. We demonstrated that factors that are known to affect multisensory integration can cause a visual task to be affected by an illusion, or remain fully unaffected, whereas a visuomotor task can be affected by an illusion, remain immune from it, or, surprisingly, even show a robust reverse effect. Thus, similar or dissociable effects can be observed as a function of factors that affect multisensory processing [30]. In addition, dissociable effects are not limited to the classic notion that action is immune to illusions, whereas perception is not [7]. To the best of our knowledge, this work is the first to document that grasping can also show inverse illusion effects in conditions whereby matching yields standard effects. These findings are relevant to attempts to resolve a long standing debate in behavioral cognitive neuroscience, in particular in meta-analyses [48-49] of this literature which have so far neglected to evaluate factors affecting multisensory processing.

In conclusion, we consider potential limitations of the current study. First, consider the stimuli, which were commercially available familiar objects. As discussed in the Stimuli section, the color and texture of our large and small inducers could no doubt facilitate accessing memorized features of these objects, including their physical size. There is evidence that familiar size can influence action programming [50]. These objects however were never grasped by the participants as they were only used as inducing objects. The test objects that participants grasped were instead nondescript rubber balls, which did not have an associated familiar standard size. In addition, an effect of familiar size would only cause the representation of the object to be more veridical. Instead, we observed robust illusion effects that deviated from veridicality. An effect of familiar size, if any, may at most have reduced these effect but this would not change our conclusions. Nonetheless, we acknowledge that it may be interesting to repeat the current study using wholly unfamiliar volumetric objects.

Second, consider our induction procedure, which involved repeatedly seeing or feeling the inducing object. Although we implemented a standardized procedure for this, we acknowledge that this procedure was managed autonomously by the participant. For this reason, there was some unavoidable random variability in its overall duration, both between trials performed by the same participant, and between participants. This may have caused the overall efficacy of induction to vary somewhat from trial to trial. However, the results and the adequate precision of our parameter estimates indicates that this potential added noise did not have much of an impact.

Third, and final, consider our proposed explanation for the complex pattern of results we observed across conditions, including cases where the illusion effect reversed.

Our explanation postulated that exposing participants to a larger or smaller object during the induction phase caused a temporary, plastic modification of the internal representation of the hand which became smaller or larger accordingly. Although this hypothesized modification parsimoniously accounts both for illusion and reversed illusion effect, as argued the Results and in the Discussion section, we acknowledge that we did not directly manipulate the internal model of the hand in our study. This could be done, for instance, by multisensory stimulation in a protocol akin to the rubber hand illusion [51]. This may be an interesting avenue for further research.

## Data availability

Raw data available in Excel Format as Supplemental Materials to this paper. Statistical analysis R scripts referred to in the text are available permanently on the Open Science Framework repository at https://osf.io/nqrf2/.

## Author contributions

Both authors contributed equally to all components of the paper.

## References

1. Ungerleider, L.G., & Mishkin, M. Two cortical visual systems. In D.J. Ingle, M.A. Goodale & R.J.W. Mansfield (Eds), The analysis of visual behavior. Cambridge, MA: MIT Press. 1982. 549–586.

2. Goodale, M. A., & Milner, A. D. Separate visual pathways for perception and action. Trends Neurosci. 1992; 15, 20e25.

3. Milner, A.D. & Goodale, M.A. Two visual systems reviewed. Neuropsychologia. 2008; 46, 774–785.

4. de Haan, E.H.F., Jackson, S.R. & Schenk, T. Where are we now with ‘What’ and ‘How’”? Cortex. 2018; 98, 1–7.

5. Borst, G., Thompson, W.L. & Kosslyn, S. M. Understanding the dorsal and ventral systems of the human cerebral cortex: Beyond dichotomies. Am. Psychol. 2011; 66, 624–632.

6. Schenk, T. & McIntosh, R.D. Do we have independent visual streams for perception and action? Cogn. Neurosci. 2010; 1, 52–78.

7. Aglioti, S., DeSouza, J.F., & Goodale, M.A. Size-contrast illusions deceive the eye but not the hand. Curr. Biol. 1995; 5, 679–685.

8. Kopiske, K.K., Bruno, N., Hesse, C., Schenk, T. & Franz, V.H. The functional subdivision of the visual brain: Is there a real illusion effect on action? A multi-lab replication study. Cortex. 2016; 79, 130–152.

9. Uccelli, S., Pisu, V., Riggio, L. & Bruno, N. The Uznadze illusion reveals similar effects of relative size on perception and action. Exp. Brain Res. 2019; 237, 953–965.

10. Bruno, N. & Pavani, F. Perception: A multisensory perspective. Oxford, UK: Oxford university press. 2018.

11. Gentilucci, M, Daprati, E, Gangitano, M. Haptic information differentially interferes with visual analysis in reaching-grasping control and in perceptual processes. Neuroreport. 1988; 9, 887–891.

12. Kritikos, A., Beresford, M & Castiello, U. Tactile interference in visually guided reach-to-grasp movements. Exp. Brain Res. 2002; 144, 1–7.

13. Pettypiece, C.E., Goodale, M.A & Culham, J.C. Integration of haptic and visual size cues in perception and action revealed through cross-modal conflict. Exp. Brain Res. 2010; 201, 863–873.

14. Haffenden, A. M., & Goodale, M. A. The effect of pictorial illusion on prehension and perception. Journal of cognitive Neuroscience. 1998; 10(1), 122–136.

15. Cesanek, E., Campagnoli, C., Taylor, J. A., & Domini, F. Does visuomotor adaptation contribute to illusion-resistant grasping?. Psychonomic bulletin & review. 2018; 25, 827–845.

16. Marteniuk, R.G., MacKenzie, C.L., Jeannerod, M., Athenes, S. & Dugas, C. Constraints on human arm movement trajectories. Can. J. Psychol. 1987; 41, 365–378.

17. Pavani, F. & Zampini, M. The role of hand size in the fake-hand illusion paradigm. Perception. 2007; 36, 1547–1554.

18. Bernardi, N.F., Marino, B.F., Maravita, A., Castelnuovo, G, Tebano, R. & Bricolo, E. Grasping in wonderland: altering the visual size of the body recalibrates the body schema. Exp. Brain Res. 2013; 226, 585–594.

19. Heed, T., Gründler, M., Rinkleib, J., Rudzik, F. H., Collins, T., Cooke, E., & O’Regan, J. K. Visual information and rubber hand embodiment differentially affect reach-to-grasp actions. Acta psychologica. 2011; 138(1), 263–271.

20. Marino, B.F., Stucchi, N., Nava, E., Haggard, P. & Maravita, A. Distorting the visual size of the hand affects hand pre-shaping during grasping. Exp. Brain Res. 2010; 202, 499–505.

21. Säfström, D., & Edin, B. B. Task requirements influence sensory integration during grasping in humans. Learning & Memory. 2004; 11(3), 356–363.

22. Fodor, J. The modularity of mind Boston, MA: MIT press. 1983.

23. Bruno, N. & Bertamini, M. Haptic perception after a change in hand size. Neuropsychologia. 2010; 48, 1853–1856.

24. Uznadze, D. The psychology of set. New York, NY: Consultants bureau. 1966.

25. Bruno, N., Garofalo, G., Daneyko, O. & Riggio, L. Visual similarity modulates visual size contrast. Acta Psychol. 2018; 188, 122–130.

26. Calzolari, E., Azañón, E., Danvers, M., Vallar, G., & Longo, M. R. Adaptation aftereffects reveal that tactile distance is a basic somatosensory feature. Proceedings of the National Academy of Sciences. 2017; 114(17), 4555–4560.

27. Uccelli, S., & Bruno, N. The effect of the Uznadze illusion is temporally dynamic in closed-loop but temporally constant in open-loop grasping. Quarterly Journal of Experimental Psychology. 2023; 17470218231206907.

28. Uccelli, S., Sacheli, L. M., & Paulesu, E. Emergent and planned interpersonal synchronization are both sensitive to ‘tempo aftereffect contagion’. Neuropsychologia. 2023; 181, 108492.

29. Smith, P. L., & Little, D. R. Small is beautiful: In defense of the small-N. Psychonomic bulletin & review. 2018; 25, 2083–2101.

30. Stein, B.E. & Meredith, M.A. The Merging of the Senses. Cambridge, MA: MIT press. 1993.

31. Lederman, S. J., & Klatzky, R. L. Hand movements: A window into haptic object recognition. Cogn. Psych. 1987; 19, 342–368.

32. Pisu, V., Uccelli, S., Riggio, L., & Bruno, N. Action preparation in grasping reveals generalization of precision between implicit and explicit motor processes. Neuropsychologia. 2020; 141, 107406.

33. Enders, C. K., & Tofighi, D. Centering predictor variables in cross-sectional multilevel models: a new look at an old issue. Psych. Meth. 2017; 12, 121.

34. Kreft, I. G., De Leeuw, J., & Aiken, L. S. The effect of different forms of centering in hierarchical linear models. Mult. Behav. Res. 1995; 30, 1–21.

35. Baayen, R. H., Davidson, D. J., & Bates, D. M. Mixed-effects modeling with crossed random effects for subjects and items. J. Mem. Lang. 2008; 59, 390–412.

36. Bates, D., Mächler, M., Bolker, B., & Walker, S. Fitting linear mixed-effects models using lme4. arXivpreprint arXiv:1406.5823. 2014.

37. Symonds, M. R., & Moussalli, A. A brief guide to model selection, multimodel inference and model averaging in behavioural ecology using Akaike’s information criterion. Behav. Ecol. Sociobiol. 2011; 65, 13–21.

38. Wasserstein, R. L., Schirm, A. L., & Lazar, N. A. Moving to a world beyond “p< 0.05”. The American Statistician. 2019; 73(sup1), 1–19.

39. Cumming, G. The new statistics: Why and how. Psych. Sci.. 2014; 25, 7–29.

40. Kline, R. B. Beyond significance testing: Reforming data analysis methods in behavioral research APA. 2004.

41. Meredith, M.A. & Stein, B.E. Interactions among converging sensory inputs in the superior culliculus. Science. 1983; 221, 389–391.

42. Holmes, N.P. & Spence, C. Multisensory integration: space, time, and superadditivity. Curr. Biol. 2005; 15, R762–R764.

43. Stein, B.E., Huneycutt, W.S. & Meredith, M.A. Neurons and behavior: the same rules of multisensory integration apply. Brain Res. 1988; 448, 355–358.

44. Spence, C. Just how important is spatial coincidence to multisensory integration? Evaluating the spatial rule. Ann NY Acad Sci. 2013; 1296, 31–49.

45. Schot, W. D., Brenner, E., & Smeets, J. B. Unusual prism adaptation reveals how grasping is controlled. Elife. 2017; 6, e21440.

46. Kopiske, K. K., Cesanek, E., Campagnoli, C., & Domini, F. Adaptation effects in grasping the Müller-Lyer illusion. Vision Research. 2017; 136, 21–31.

47. Harris, C. S. (1963). Adaptation to displaced vision: visual, motor, or proprioceptive change?. Science. 1963; 140(3568), 812–813.

48. Franz, V.H. & Gegenfurtner, K.R. Grasping visual illusions: consistent data and no dissociation. Cogn. Neuropsychol. 2008; 25, 920–950.

49. Bruno, N. & Franz, V.H. When is grasping affected by the Müller-Lyer illusion? A quantitative review. Neuropsychologia. 2009; 47, 1421–1433.

50. McIntosh, R. D. & Lashley, G. Matching boxes: familiar size influences action programming. Neuropsychologia. 2008; 46, 2441–2444.

51. Botvinick, M., & Cohen, J. Rubber hands ‘feel’ touch that eyes see. Nature. 1998; 391(6669), 756.

